# Single-neuron interactions between the somatosensory thalamo-cortical circuits during perception

**DOI:** 10.1101/315911

**Authors:** Adrià Tauste Campo, Yuriria Vázquez, Manuel Álvarez, Antonio Zainos, Román Rossi-Pool, Gustavo Deco, Ranulfo Romo

## Abstract

Sensory thalamo-cortical interactions are key components of the neuronal chains associated with stimulus perception, but surprisingly, they are poorly understood. We addressed this problem by evaluating a directional measure between simultaneously recorded neurons from somatosensory thalamus (VPL) and somatosensory cortex (S1) sharing the same cutaneous receptive field, while monkeys judged the presence or absence of a tactile stimulus. During the stimulus-presence, feedforward (VPL→S1) interactions increased, while pure feedback (S1→VPL) interactions were unaffected. Remarkably, bidirectional interactions (VPL↔S1) emerged with high stimulus amplitude, establishing a functional thalamo-cortical loop. Furthermore, feedforward interactions were modulated by task context and error trials. Additionally, significant stimulus modulations were found on intra-cortical (S1→S1) interactions, but not on intra-thalamic (VPL→VPL) interactions. Thus, these results show the directionality of the information flow between the thalamo-cortical circuits during tactile perception. We suggest that these interactions may contribute to stimulus perception during the detection task used here.

## INTRODUCTION

A major challenge in systems neuroscience involves understanding how perceptual experiences arise from coordinated neural interactions, and how information flows among the interacting neurons (Salinas et al., 2000; de Lafuente and Romo, 2005; Hernández et al., 2010; Tauste Campo et al., 2015). The sensory thalamus is an essential node within the perceptual circuit, relaying information from the periphery to the cortex (Sherman, 2016). Given its connectivity with the cortex, feedback cortical inputs can favor information transmission (Alitto and Usrey, 2003; Briggs and Usrey, 2007; Crandall et al., 2015; Wang et al., 2007). Thus, the thalamus could act as a processing unit in continuous interaction with the cortex (Jones, 2002; Sherman, 2016; Sherman and Guillery, 2006). Evidence supporting this view relies from previous studies in anesthetized (Ahissar et al., 2000; Alonso et al., 1996; Bruno and Sakman, 2006) and awake animals (Bruno, 2011; Wang et al., 2007). However, it is unclear how feedforward and feedback interactions within the thalamo-cortical circuit co-exist and if they correlate with the subject’s perception.

To address these questions, we recorded the simultaneous activity of single neurons in the ventral posterolateral nucleus (VPL) of the somatosensory thalamus and in primary somatosensory cortex (S1) sharing the same cutaneous receptive field, while monkeys performed a vibrotactile detection task. The animals were trained to report the presence or absence of a tactile stimulus of variable amplitude. In this task, previous works showed that VPL and S1 neurons encode mostly the physical features of the stimulus (Vázquez et al., 2012, 2013). These findings rise several questions. First, how sensory information is communicated between the VPL and S1 at the level of single neurons during the stimulus relay and perception? Second, what is the balance between the information flowing in a feedforward direction (VPL→S1) and in a feedback (S1→VPL) direction? Third, does the information flow between VPL and S1 correlate with the subject’s perception and behavioral context?

In the current work, we addressed the above questions by detecting time-varying directional couplings between the recorded VPL and S1 neuron pairs across many trials during the vibrotactile detection task. Our results reveal that, in the presence of a tactile stimulus, feedforward (VPL→S1) interactions largely prevail over feedback (S1→VPL) interactions. Interestingly, bidirectional interactions (VPL↔S1) emerged in a non-negligible number, which were modulated by the stimulus amplitude above detection threshold. Importantly, bidirectional interactions constitute a novel mechanism that could allow synergic cooperation between these areas to transmit stimulus information. Additionally, neurons that were involved in bidirectional interactions exhibited higher firing rates in both areas. Critically, at the stimulus onset, feedforward interactions correlated with the subject’s perception. During a variant of the detection task (passive condition, in which no response was required from the animal), feedforward interactions were reduced during the possible window of stimulation for both the stimulus-present and stimulus-absent trials, while bidirectional interactions were only reduced in the second half of the stimulus. Finally, VPL-S1 interactions were contrasted against intra-area interactions. Notably, during the stimulation epoch, S1-S1 interactions increased in both types: unidirectional and bidirectional. In contrast, VPL-VPL interactions were barely affected with the arrival of the stimulus. Thus, our results, characterize the directionality of information flow between VPL-S1 neuron pairs during tactile perception. They reveal that feedforward interactions besides relaying stimulus information, convey task-context information, whereas bidirectional interactions might reflect coordinated cooperation between VPL and S1 to transmit information about the stimulus, when it is detected above the behavioral threshold.

## RESULTS

Two monkeys (*Macaca Mulatta*) were trained to perform a tactile detection task (de Lafuente and Romo, 2005 and 2006). In each trial, the animal had to report whether the tip of a mechanical stimulator vibrated or not (Fig. 1A). Stimuli were sinusoidal, had a fixed frequency of 20 Hz and were delivered to the glabrous skin of one fingertip; crucially, they varied in amplitude across trials. Stimulus-present trials were interleaved with an equal number of stimulus-absent trials in which no mechanical vibrations were delivered (Fig. 1A). The presence or absence of the stimulus (0.5 s) was preceded by a variable pre-stimulus period (1.5–3 s), followed by a fixed post-stimulus delay period of 3 s before the monkey reported its decision by pressing one of two push-buttons (Fig. 1A). Stimulus detection thresholds were calculated from the behavioral responses (left panel of Fig. 1B). Importantly, depending on the monkeys’ responses, trials could be classified into four types: hits and misses in the stimulus-present trials, and correct rejections and false alarms in the stimulus-absent condition (right panel of Fig. 1B). Once the animals performed the task at near detection threshold (8 μm), we recorded the simultaneous activity of spike trains from individual neurons from VPL and S1 (areas 3b or 1, Fig. 1C), while monkeys performed the task. Here, it is important to highlight that the recorded neuron pairs (n = 84) from the VPL and S1 shared exactly the same cutaneous receptive fields (Fig. 1D). Importantly, one or more than two neurons from VPL and S1 were simultaneously recorded in the same session. This means that we can use the same dataset to study VPL-S1 and intra-areas interactions or the response population of each area. Fig. 1E shows that the neuronal populations of VPL and S1 modulate their firing rate during the stimulus-present condition (left panel), but not during the stimulus-absent condition (right panel). Thus, it appears an optimal experimental condition for assessing the time-varying directionality of information flow between neuron pairs from VPL and S1 during the detection task.

**Figure 1.**
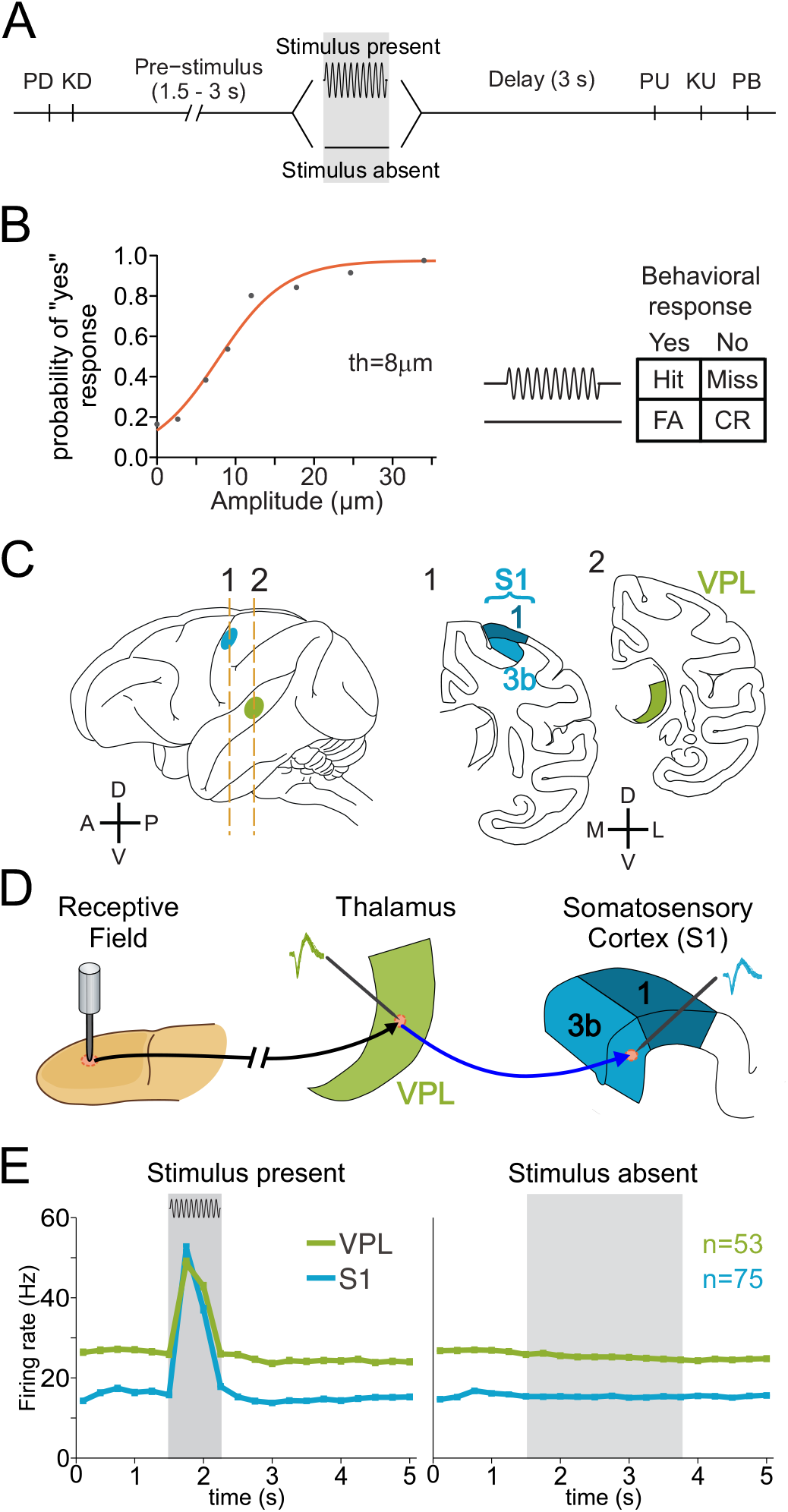
Detection task, psychophysical performance, recording sites and neuronal responses during the task. (A) Vibrotactile detection task. Trials began when the stimulator probe indented the skin of one fingertip of the monkey’s restrained right hand (probe down, PD); the monkey reacted by placing its left, free hand on an immovable key (key down, KD). After a variable pre-stimulus period (1.5 – 3 s), a vibratory stimulus of variable amplitude (1−34 μm, 20 Hz, 0.5 s duration) was presented on one half of the trials; no stimulus was presented on the other half of the trials. Following the stimulus presentation period or a period where no stimulus was delivered, the monkey waited for 3 s until the probe was lifted off from the skin (PU, probe up); then the animal removed its free hand from the key (KU, key up) and pressed one of two push buttons (PBs) to report whether the stimulus was present or absent. Lateral and medial buttons were used for reporting stimulus-presence and stimulus-absence, respectively. Stimulus-present and stimulus-absent trials were randomly interleaved within a run. (B) The button pressed indicated whether the monkey felt the stimulus (henceforth referred as ‘yes’ and ‘no’ responses, respectively). Left panel in B: mean psychometric function depicting the probability of the monkey reporting ‘yes’ as a function of the stimulus amplitude (th = 8 μm, detection threshold). Right panel in B: behavioral responses depending on the stimulus-presence (Hit or Miss) or stimulus-absence (CR, correct rejection; FA, false alarm). (C) Recording sites in the ventral posterior lateral (VPL) nucleus (green) of the thalamus and in areas 1 and 3b of the primary somatosensory cortex (S1, cyan). (D) Scheme depicting how the neural activity from single neurons in the VPL and S1 (3b or area 1) sharing the same cutaneous receptive field was simultaneously recorded during the detection task. (E) Mean firing rate for the simultaneously recorded VPL (n = 53) and S1 (n = 75) neurons during the stimulus-present and stimulus-absent trials.

### Assessing directional interactions between VPL and S1

It is well established that the VPL relays information from the skin mechanoreceptors up to S1 (Fig. 1C and 1D). This knowledge allows us to test the following hypothesis in the detection task: after the stimulus onset the amount of directional interactions becomes crucially larger in the VPL→S1 direction than in the S1→VPL direction. However, given the larger connectivity from S1 to VPL than from VPL to S1 (Sherman and Guillery, 2006), the second hypothesis is that the amount of direction information flow is higher from S1→VPL than from VPL→S1. Importantly, we also hypothesize that the direction information flow between VPL and S1 could be affected by the task conditions (Fig. 1B). We addressed all these hypotheses by using a non-parametric method that measures direct information flow between the simultaneously recorded spike trains of pairs of VPL-S1 neurons in single trials and within sequential time windows (Massey 1990, Jiao et al., 2013, Tauste Campo et al., 2015, STAR Methods). The method is illustrated in Fig. 2A. For every pair of simultaneous spike trains from VPL-S1 neurons, we estimated the directed information (DI) conveyed in both directions (VPL→S1 and S1→VPL) at time delays of [0,2,4…,20]ms, in non-overlapping and consecutive time windows of 0.25 s along each trial during the detection task (Fig. 2A, left panel). To infer the significance of each estimation, we defined a maximizing-delay statistic (*T*) and built the corresponding null distribution (〈T〉) using circular-shifts of the target sequences (Fig. 2A, middle panel). For each (single-trial) directional pair (X → Y), the method assessed the significance of the interaction together with an unbiased estimation of the statistic value and the maximizing-delay (Fig. 2A, right panel). Directional pairs associated with significant estimators (*α* = 0.05) will be next referred as directional interactions. To address statistical differences between the distinct types of interactions (e.g. VPL→S1 versus S1→VPL) at every time interval, we performed a non-parametric paired test for correlated samples (Winkler et al, 2014) with FWER (family wise error rate) significance level *α* = 0.05 (STAR Methods). In addition, we quantified Cohen’s h effect size for proportion differences (H, Cohen, 1988)) to control for the influence of large sample size (> 100 samples) into significant findings. Therefore, for the percentage of directional interactions, we first underlined those consecutive time intervals where the difference was significant (P < 0.05, FWER) and the effect size was larger than H = 0.3 (in grey) and H = 0.5 (in cyan), respectively (STAR Methods).

**Figure 2.**
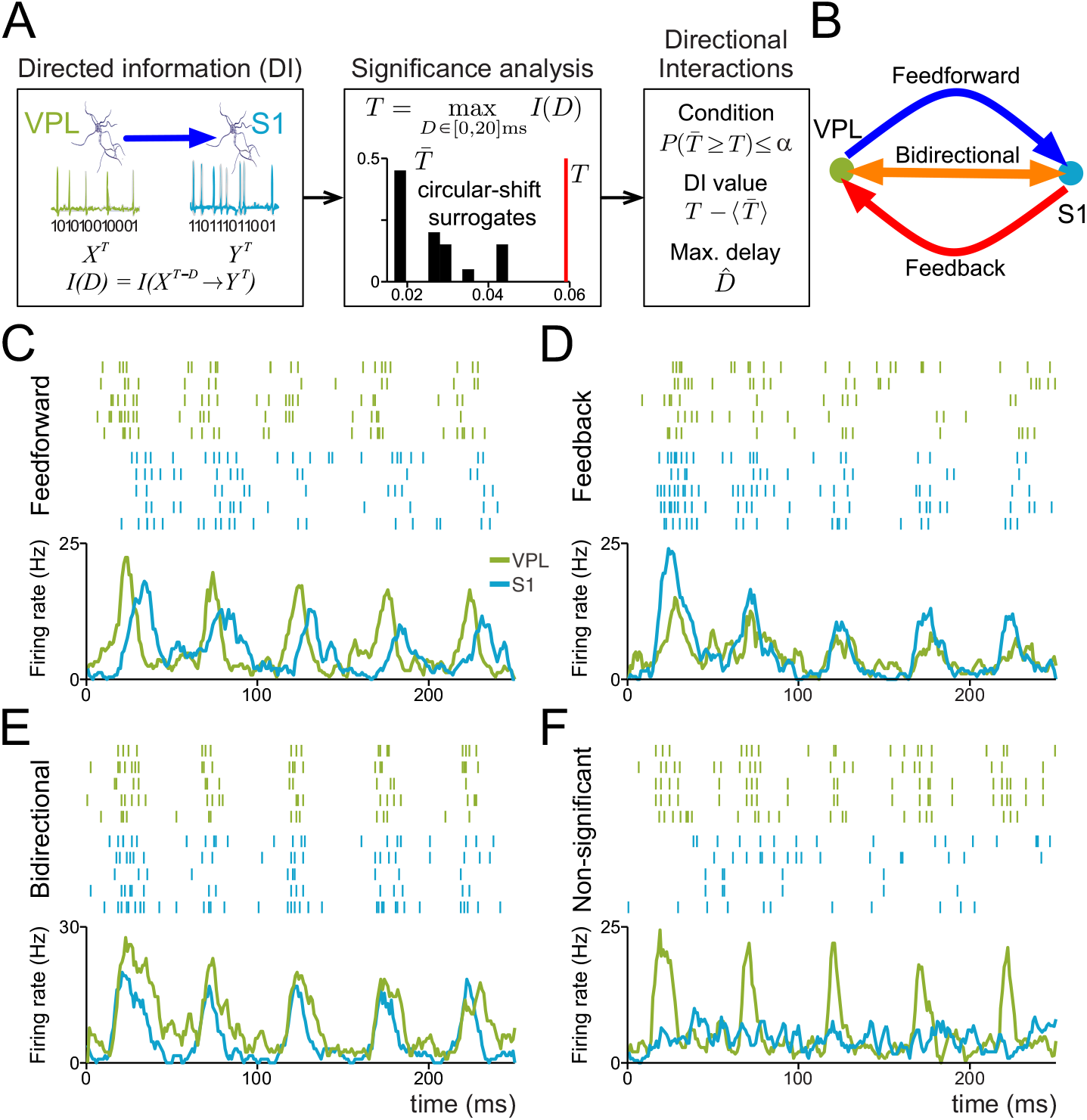
Assessing directional interactions between pairs of VPL and S1 neurons during the detection task. (A) Sequential scheme representing the method to infer single-trial directional interactions. Left panel: directed information is estimated between single-trial spike trains of the simultaneously recorded neurons in VPL and S1 for delays (0, 2,…, 20) ms. Middle panel: significance is locally determined via non-parametric testing (α = 0.05). Right panel: every significant interaction is associated with an unbiased directed information value (T-〈T〉) and a maximizing delay (D̂). (B) Graphical representation for feedforward (VPL→S1, in blue), feedback (S1→VPL, in red) and bidirectional (S1↔VPL, in orange) interactions between VPL and S1 neurons. (C-F) Raster plots and spike density functions depicting the neural activity occurring during the first 250 ms of stimulation (34 μm) for examples of VPL (green) and S1 (cyan) neuron pairs involved in feedforward (C), feedback (D), bidirectional (E), and non-significant (F) directional interactions.

A first characterization of the VPL-S1 interactions was done by measuring the percentage of directional interactions (Fig. S1A), as well as the mean directed information value (Fig. S1B) for: VPL→S1 and S1→VPL directions during the time course of the detection task. Both quantities were calculated separately for the stimulus-present (left) and stimulus-absent (right) trials. This analysis showed that for the stimulus-present trials, both the number of the VPL→S1 interactions (trials = 3216 in neuron pairs = 84, P < 0.01, H > 0.5, cyan line; Fig. S1A left) and the mean directed information value (P < 0.01, D > 0.15, gray lines; Fig. S1A left) were significantly larger than the S1→VPL interactions during the first half (250 ms) of the stimulus period and weaker, but still significant during the second half (H < 0.5; D > 0.15). In contrast, for stimulus-absent trials, the greatest differences were manifested by the number of interactions (trials = 4371 in neuron pairs = 84, P < 0.01, H > 0.3; Fig. S1 right) during the possible stimulation window (PSW, 1.5-3.5 s) with no counterpart evidence from the mean directed information value. Along the text, we will refer to the possible stimulation window (PSW) as the temporal window within a trial when the monkey was expecting a stimulus (Carnevale et al., 2015). We restricted our analysis to the amount of directional interactions (rather than their average value) due to its greater sensitivity to detect directionality differences during the stimulus and possible stimulation windows.

As a first approach, we considered exclusively two directionality cases: VPL→S1 and S1→VPL (Fig. S1). Yet, when studying interactions for two simultaneous spike trains *X* and *Y* at a given time interval (e.g., 250 ms) one may instead consider three cases: the spike trains are coupled in only one direction (*X* → *Y*), in the opposite direction (*Y* → *X*) or simultaneously coupled in both directions (*X* ↔ *Y*). In principle, these three cases correspond to neurons in each pair taking three different roles: driver, target or both, which may be associated with distinct functional mechanisms (Deschênes et al., 1998). According to this notion, we classified each directional interaction by pairing the location and role of each neuron per trial. In short, we defined as feedforward interactions (VPL→ S1) those interactions involving pairs where the neuron of VPL was the driver and the neuron of S1 was the target. Similarly, we defined as feedback interactions (S1→VPL) where the neuron of S1 was the driver and the VPL neuron was the target. Finally, pairs where neurons exhibited simultaneously feedforward and feedback interactions were labeled as bidirectional interactions. Figure 2B shows a schematic representation for the three types of interactions and Fig. 2C-E shows four example pairs of neurons (five repetitive trials; green color for VPL raster plots and cyan color for S1 raster plots) responding during the first 250ms after stimulus onset with significant feedforward (Fig. 2C), feedback (Fig. 2D) and bidirectional (Fig. 2E) interactions, together with non-significant interactions (Fig. 2F).

### Stimulus-presence modulates feedforward and bidirectional interactions

Having set the above definitions, we first examined for each type of interaction the mean interaction delay as a putative intrinsic property of each interaction (Fig. S2). Bidirectional interactions occurred at shorter delays than unidirectional interactions for both stimulus-present (trials = 3217 in neuron pairs = 84, 〈D = 0.35〉, time-average Cohen’s *d*; Fig. S2A left) and stimulus-absent trials (trials = 4731 in neuron pairs = 84, 〈D = 0.37〉, time-average Cohen’s *d*; Fig. S2A right). The delay histograms (Fig. S2B) revealed that these differences were, in general, due to higher percentage of zero-delay couplings (around 40%) in bidirectional interactions as compared with unidirectional interactions (around 15%). Further, we examined the interaction delay distribution during the stimulus-present trials. (Fig. S2B, top-left panel). We found that the arrival of the stimulus increased the relative amount of bidirectional interactions at zero delay (from 36% to 49%, Fig. S2B, top-left panel) and the relative amount of feedforward interactions at 8 ms (from 9% to 15%), while feedback interactions were only moderately enhanced at 8 ms (from 9% to 11%).

Second, we investigated two main questions regarding feedforward, feedback and bidirectional interactions: 1) what was the contribution of each interaction type to a greater number of VPL→S1 interactions observed during the stimulus-presence? 2) What was the contribution of each neuron pair to this effect? Figure 3A depicts the percentages of directional interactions according to their type (feedforward, feedback, and bidirectional) for stimulus-present (left) and stimulus-absent trials (right) during the time-course of the task. The proposed decomposition highlighted the contribution of each type into the directionality differences illustrated in Fig. S1. The increment in the VPL→S1 direction, especially after the stimulus onset, was mainly contributed by feedforward interactions (trials = 3216 in neuron pairs = 84, P < 0.05, H = 0.23; Fig. 3A left, blue trace) and lesser by the bidirectional interactions (trials = 3216, neuron pairs = 84, P < 0.05, H = 0.1; Fig. 3A left, orange trace). In contrast, the increase in the S1→VPL direction (Fig. S1) could only be explained by an increase in bidirectional interactions. Indeed, genuine feedback interactions were not significantly modulated during the stimulus-presence (P > 0.05; Fig. 3A left, red trace). On the other hand, for stimulus-absent trials, none interaction type showed a significant modulation (trials = 4271 in neuron pairs = 84, P > 0.05; Fig. 3A right).

**Figure 3.**
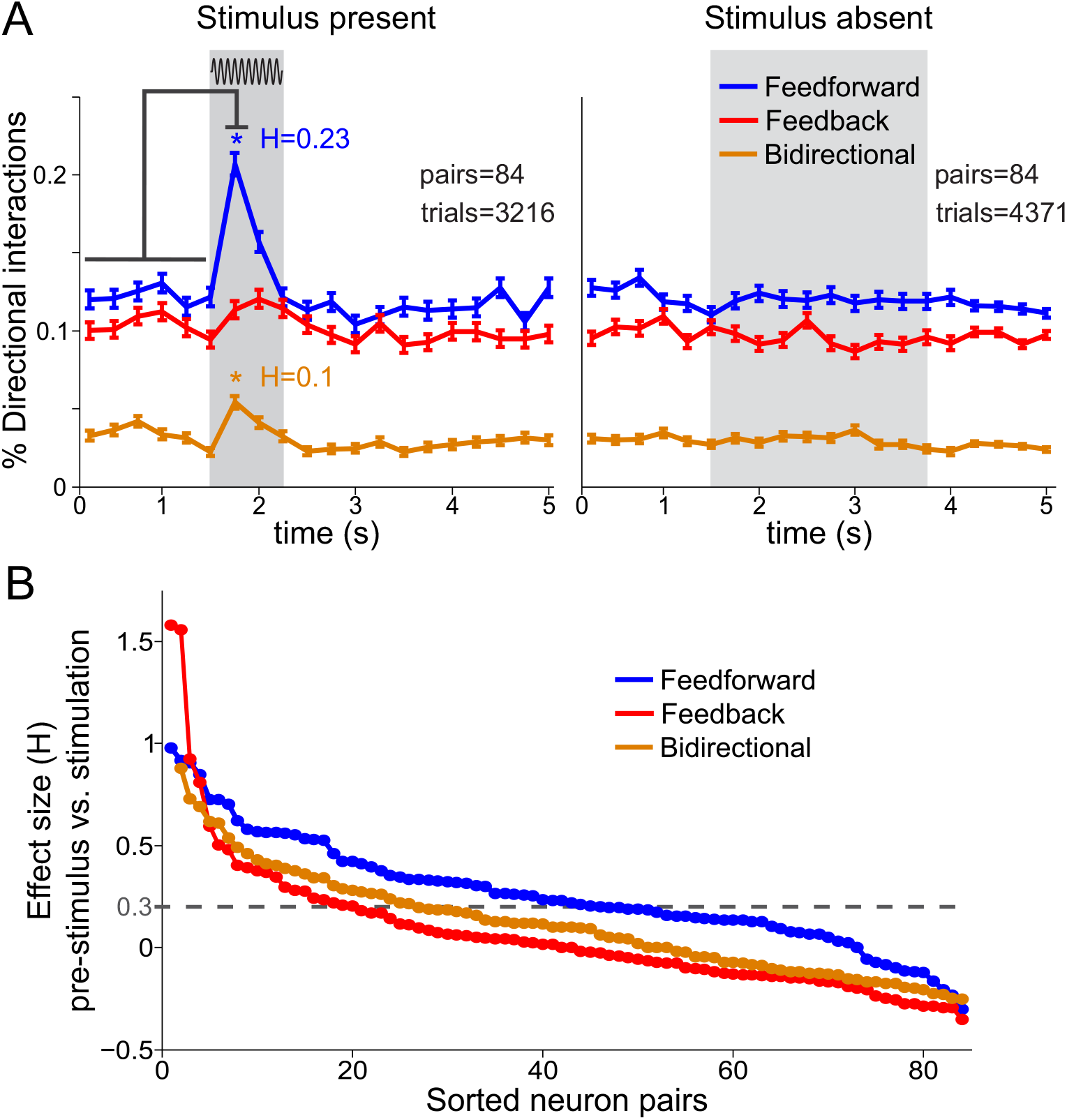
VPL-S1 feedforward, feedback and bidirectional interactions during the detection task. (A) Percentage of feedforward, feedback and bidirectional interactions during the time course of the task during the stimulus-present (left panel, trials = 3216 hits; neuron pairs = 84) and stimulus-absent trials (right panel, trials = 4371 correct rejections; neuron pairs = 84). Error bars denote the SEM (standard error of the mean). In all figures, gray boxes depict the stimulation period for the stimulus-present trials, and the possible window of stimulation (PWS) for the stimulus-absent trials. Asterisks denote significant differences (P < 0.05) between the pre-stimulus and the first stimulation interval (non-parametric test, α = 0.05). H denotes the effect size and average effect size (Cohen’s H) of significant differences. (B) Effect size of the difference between the percentage of each interaction type before the stimulus period (first 6 task intervals, 1.5 s) and during the first stimulation interval (0-0.25 s) as a function of all neuron pairs for distinct directional interactions. Each plot is represented by ordering the neuron pairs in descending order according to the magnitude of the effect size for each particular interaction type. The black horizontal dashed line represents the effect size threshold (H ≥ 0.3).

We then investigated the contribution of each neuronal pair to the increment in feedforward and bidirectional interactions during the first half of the stimulation period in the stimulus-present trials. To do so, we repeated the analysis of Fig. 3A for every VPL-S1 pair and obtained a stimulus-driven effect size per pair. Figure 3B illustrates these results by sorting the neuronal pairs in a descending effect size order. Consistent with our previous findings, the stimulus effect was larger in feedforward and bidirectional interactions than in feedback interactions in most of the neuronal pairs. Besides that, approximately 40% (33/84) and 20% (18/84) of the neuronal pairs increased the number of feedforward and bidirectional interactions, respectively, with moderate and large effect sizes (H > 0.3, Fig. 3B).

Next, we asked how often a VPL-S1 pair could simultaneously handle both interaction modulations during the stimulation period. To address this question, we correlated the stimulus effect sizes associated with each type of interaction across all the recorded VPL-S1 pairs. The correlation value between the effect sizes was rather low (ρ = −0.11, P > 0.5, Spearman’s rho), suggesting that feedforward and bidirectional VPL-S1 interactions might be modulated by a different thalamo-cortical mechanism. We also analyzed how neuronal pairs were differently modulated during stimulus-absent trials (Fig. S3). To this end, we compared the percentage of feedforward, feedback and bidirectional interactions per neuronal pair during the first interval of the stimulus window and during the first interval of the PSW. Each point in Fig. S3, represents a VPL-S1 pair and the angular distribution of these points are shown as insets. Therefore, angular values smaller than 45° indicated stronger interaction during the stimulus period. Thus, mean angular values indicated that the stimulus-present effect (θ < 45°) was more prominently manifested in feedforward (θ = 30.5°; Fig. S3) and bidirectional interactions (θ = 25.2°; Fig. S3) than in feedback interactions (θ = 41.1°; Fig. S3). In sum, we provided consistent evidences from independent analyses that tactile stimuli mainly modulated feedforward and bidirectional interactions in VPL-S1 pairs.

### Correlation between the firing rate and the type of directional interaction

By construction, the statistical method used in this study infers interaction measurements that are not biased by the fluctuations of the neuronal spikes. To empirically corroborate this fact, we examined the influence of the firing rate into the observed VPL-S1 directional interactions. In previous related works (Vázquez et al., 2012 and 2013), we showed that the average firing rate was larger in VPL than in S1 neurons attaining similar values during the stimulation period, as shown here (Fig. 1E). Therefore, the reported increase of feedforward interactions (as compared to feedback) occurred while the VPL and S1 neurons exhibited similar firing rates. This initially suggested a low dependence between the firing rate and the existence of directional interactions. We then tested this hypothesis by correlating the firing rate driver and target neurons with the existence of incoming/outgoing directional interactions. The obtained Spearman correlation coefficients were significant, but rather low for both driver (intervals = 206360 in neuron pairs = 84, ρ = 0.11, P < 0.05; Fig. S4) and target neurons (intervals = 206360 in neuron pairs = 84, ρ = 0.07, P < 0.05; Fig. S4A). Moreover, these results were stable during stimulus-absent trials (intervals = 206280 in neuron pairs = 84, ρ = 0.1, ρ = 0.07, P < 0.05; Fig. S4B). This shows that the increase in the number of directional interactions during the stimulus period could not be merely explained by an increase in the mean firing rate of either the VPL or S1 neurons.

The above results suggested that the firing rate might be poorly correlated with the distinct directional interactions. To specifically address this question, we examined the firing rate of VPL and S1 neurons associated with feedforward, feedback and bidirectional interactions (Fig. 4). Our analysis revealed that the firing rate of VPL and S1 neurons was not significantly different (n = 53 and n =75, respectively, P > 0.05; Fig. 4) between feedforward and feedback interactions across all task intervals, including those from the stimulation period. In contrast, neurons holding bidirectional interactions showed a significant increase in their firing rate with respect to unidirectional interactions during the first interval of the stimulus period (P < 0.05; Fig. 4). These increases were manifested at supra-threshold amplitudes for VPL neurons (n = 53, P < 0.001; Fig. 4) during the entire stimulus period, while for S1 neurons (n = 75, P < 0.05; Fig. 4) only during the first interval of stimulation. Interestingly, for stimulus-absent trials, bidirectional neurons showed enhanced firing rates during PSW intervals as compared to unidirectional neurons. In sum, these results show that firing rates were not able to discriminate between unidirectional interactions (feedforward vs. feedback), but could be significantly higher for bidirectional interactions during specific task intervals (stimulus onset or PSW). Importantly, this refined firing-rate analysis provides initial evidence that bidirectional neurons might be task-modulated regardless of the stimulus-presence.

**Figure 4:**
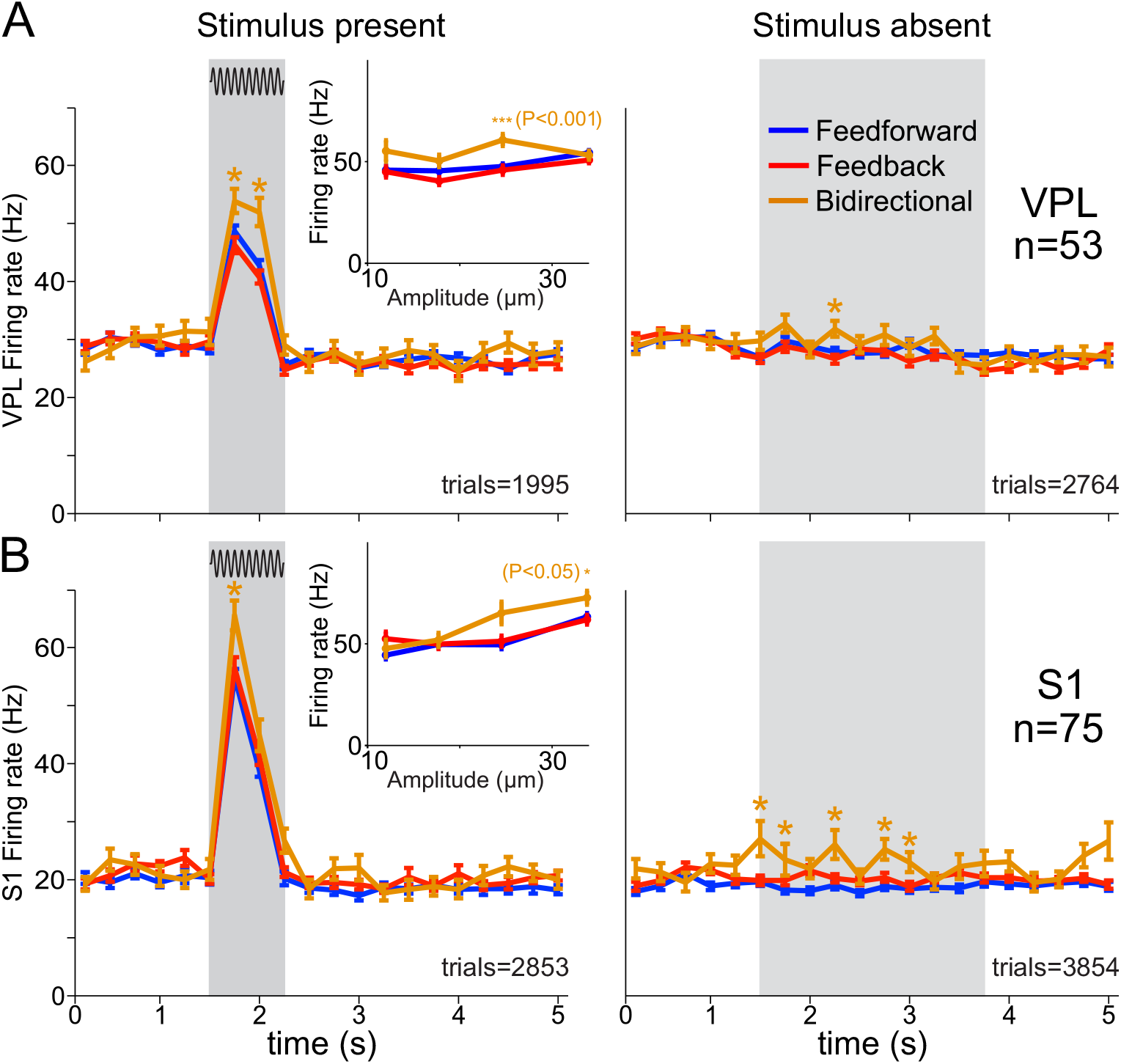
Mean firing rate of VPL and S1 neurons during detection task for feedforward, feedback and bidirectional interactions. For all panels, gray boxes depict the stimulation period for the stimulus-present trials (left panels), and the possible window of stimulation (PWS) for the stimulus-absent trials (right panels). Blue, red and orange traces depict the mean firing rate for neurons holding feedforward, feedback and bidirectional interactions, respectively, during the time course of the detection task (neuron pairs = 84). Error bars denote the SEM (standard error of the mean). (A) Mean firing rate for VPL neurons (n = 53; left panel, trials = 1995 hits; right panel, trials = 2764 correct rejections). Insets depict the mean firing rate as a function of the stimulus amplitude. Asterisks denote significant differences (*, P<0.05) among the firing rate of neurons holding distinct directional interactions. (B) Similar as panel A, but for S1 neurons (n = 75; left panel, trials = 2853 hits; right panel, trials = 3854 correct rejections).

### Directional interactions within and across VPL and S1 areas

So far, we have tested the influence of the local firing rates into the emergence of interactions across the VPL and S1. However, it is unknown whether the local interactions within each of these areas contribute to the observed interactions across areas. In other words, what are the roles of these interactions within the VPL or S1? To address this question, we repeated again our directionality analysis over all simultaneously recorded neuronal pairs within the VPL and within S1, separately (Fig. 5). In each area, the terms feedforward and feedback were substituted by arbitrary unidirectional 1 and unidirectional 2 interactions across pairs within the same area. Although several realizations were possible under these arbitrary choices, Fig. 5 depicts a representative realization for each area together with the VPL-S1 results reported in Fig. 3. While the number of simultaneous pairs in S1 still led to a large number of stimulus-present trials (trials = 1650 in neuron pairs = 42; Fig. 5A) and stimulus-absent trials (trials = 2244 in neuron pairs = 42; Fig. 5B), the low number of VPL trials collected in few pairs (trials = 234 in neuron pairs = 6; Fig. 5) notably limited the study of differences in VPL-S1 interactions. Yet, contrary to VPL-S1 pairs, the results showed that there were no sustained significant differences in S1 and VPL intra-area unidirectional interactions in both experimental conditions (P > 0.05; Fig. 5). Additionally, the three types of S1-S1 interactions showed a significant increase during the stimulation period, which were more sustained over stimulus-presence than for VPL-S1 pairs. This result supports the idea that S1 neurons coordinate their activity during the entire stimulation period. In contrast to this result, the VPL-VPL interactions did not show any modulation during the stimulus-presence.

**Figure 5.**
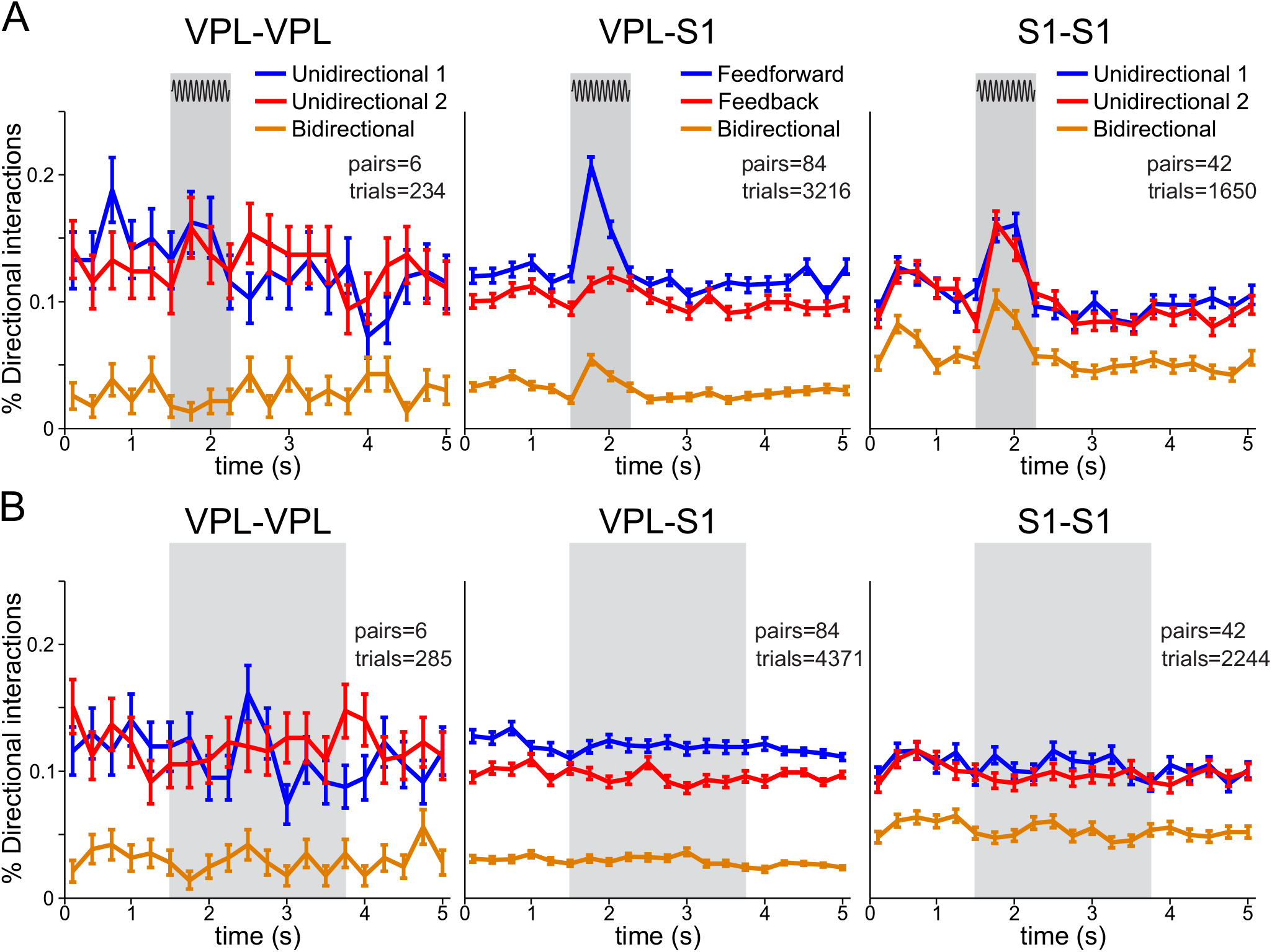
Directional interactions during the detection task for VPL-VPL, S1-S1 and VPL-S1 neurons. (A and B) Left panels depict the percentage of interactions in one arbitrary direction (blue), the opposite (red) and both (orange) directions within VPL neurons (trials = 234 hits; trials = 285 correct rejections; neuron pairs = 6). (A and B) Middle panels depict the percentage of feedforward (blue), feedback (red) and bidirectional (orange) interactions across VPL-S1 neurons (trials = 3216 hits; trials = 4371 correct rejections, neuron pairs = 84). (A and B) Right panels depict the percentage of interactions in one arbitrary direction (blue), the opposite (red) and both (orange) within S1 neurons (trials = 1650 hits; trials = 2244 correct rejections; neuron pairs = 42). Error bars denote the SEM (standard error of the mean). (A) Time course of directional interactions during the stimulus-present trials. (B) Time course of directional interactions during stimulus-absent trials.

### The amount of VPL-S1 interactions is modulated by the stimulus amplitude

We showed above that feedforward and bidirectional VPL-S1 interactions were enhanced by the stimulus-presence, while feedback interactions were minimally affected (Fig. 3). Furthermore, each interaction was modulated by different neuronal populations. This prompted to hypothesize that feedforward and bidirectional interactions could be differently related to the stimulus amplitude. To further investigate this question, we divided the stimulus-present trials into 3 groups based on the stimulus amplitude. We grouped the 9 μm trials within the near-threshold (middle left panel of Fig. 6) group and defined the supra-threshold (left most panel of Fig. 6) and sub-threshold (middle right panel of Fig. 6) groups as those stimulus-present trials above and below 9 μm, respectively. We choose 9 μm as the cut-off amplitude since it is the closer amplitude to the monkey’s mean detection threshold (8 μm; Fig. 1B). We also considered stimulus-absent trials for comparison (right most panel of Fig. 6A). First, we quantified the amount of interactions within each group (Fig. 6A). Notably, we found that the stimulus effect observed for all amplitudes (Fig. 3B) was mainly due to the supra-threshold group (> 9μm). At these amplitudes, feedforward and bidirectional interactions showed a significant incremental effect (trials = 2237 in neuron pairs = 84, P < 0.01, H = 0.26 and H =0.14; Fig. 6A), while feedback interactions were not significantly altered (P > 0.05). In contrast, for the near-threshold group, the feedforward increment was preserved (trials = 443 in neuron pairs = 84, P < 0.01, H = 0.2; Fig. 6A), while bidirectional interactions dropped dramatically (P > 0.05). Finally, the sub-threshold amplitudes only showed a weaker significant increase in the amount of feedforward interactions (trials = 536 in neuron pairs = 84, P < 0.01, H = 0.13; Fig. 6A). Thus, the results illustrated in Fig. 6A demonstrated that the amount of directional interactions was amplitude dependent. To examine this dependency, we focused on the first 250 ms of the stimulus-present period and plotted the percentage of interaction types as a function of the stimulus amplitude (Fig. 6B). Besides, we computed single-trial (r) and average-trial (R) Spearman’s rho correlations between the amount of interactions and amplitude values (STAR Methods). During the first 250 ms the amount of each type of interaction was correlated with the amplitude values as measured by single-trial types (P < 0.05; Fig. 6B). But, when considering average-trial correlations, the significance analysis provided different outcomes across interaction types. Indeed, Fig. 6B shows that feedforward (trials = 7587, r = 0.12 in neuron pairs = 84, R=0.97, P <0.01; Fig. 6B) and bidirectional (trials = 7587, r = 0.06 in neuron pairs = 84, R=0.85, P < 0.05; Fig. 6B) interactions exhibited a significant monotonic modulation, while feedback interactions remained approximately constant over the amplitude values (trials = 7587, r = 0.03 in neuron pairs = 84, R=-0.24, P > 0.05; Fig. 6B). These results complemented the analysis of Fig. 6A, revealing that the amount of feedforward and bidirectional interactions was linearly associated with the stimulus amplitude. Overall, feedforward and bidirectional interaction were enhanced by the stimulus-presence and they could convey information about the stimulus amplitude.

**Figure 6.**
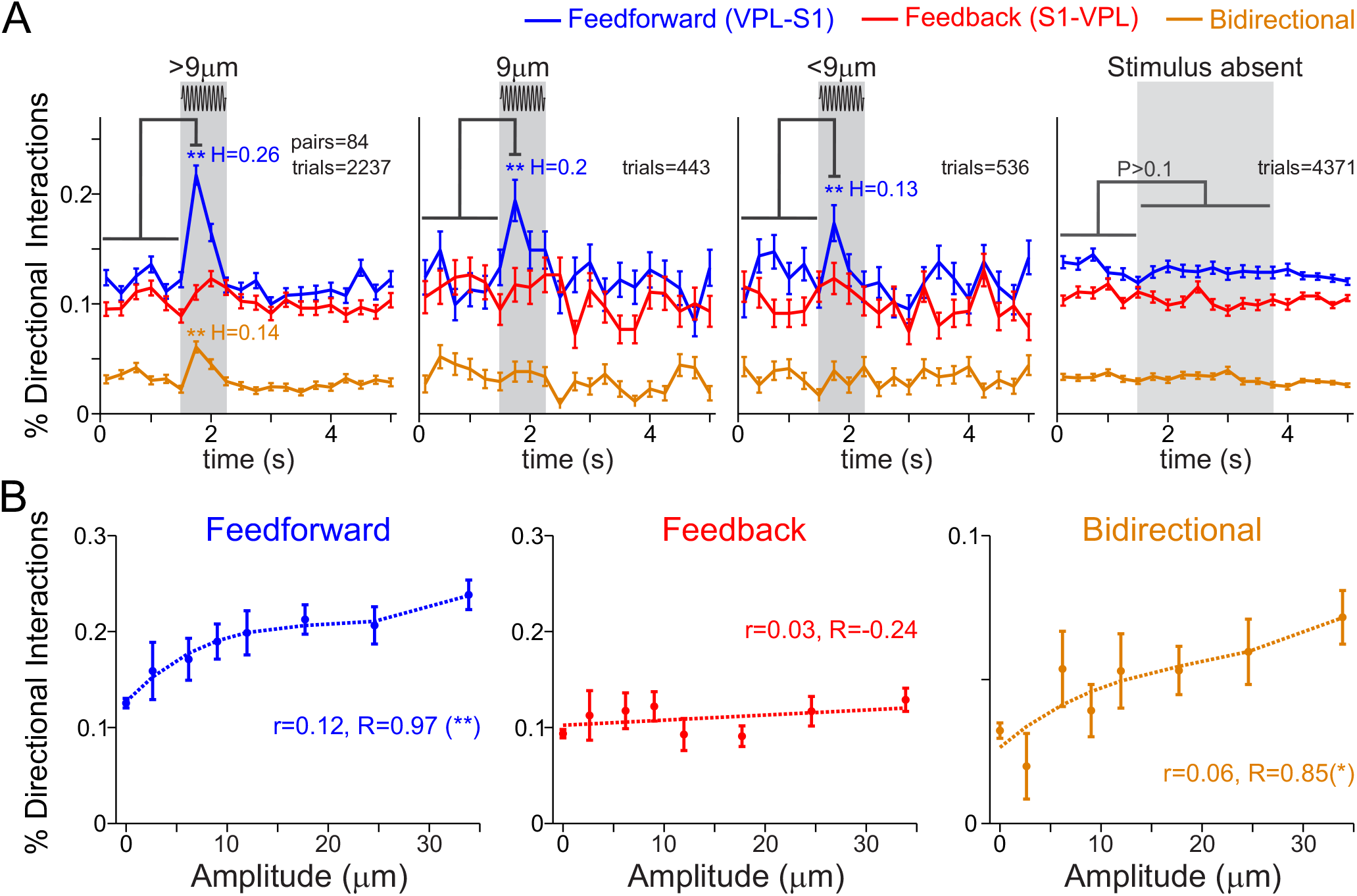
VPL-S1 feedforward and bidirectional interactions are modulated by the stimulus amplitude. (A) Time course of the percentage of feedforward (VPL→S1, blue), feedback (S1→VPL, red) and bidirectional (S1↔VPL, orange) interactions between VPL and S1 neurons during stimulus-present and stimulus-absent trials. From left to right: percentage of interactions for supra-threshold (> 9 μm, neuron pairs = 84; trials = 2237 hits), threshold (9 μm, neuron pairs = 84; trials = 443 hits), subthreshold (< 9 μm, neuron pairs = 84; trials = 536 hits) stimulus amplitudes and stimulus-absent trials (neuron pairs = 84; trials=4371 correct rejections). Asterisks denote significance levels (*, P < 0.05, **, P < 0.01). H denotes the effect size (Cohen’s h) of significant percentage differences. Error bars denote the SEM (standard error of the mean). (B) Mean percentage of feedforward, feedback and bidirectional interactions as a function of the stimulus amplitude during the first half of the stimulation period (left, 0 −0.25 s). The value of r is the correlation between the stimulus amplitude and the existence of directional interactions in each type across all trials (no amplitude-averages) with Spearman correlation (trials = 7587). The value R is the analogous correlation considering amplitude-average values (amplitudes = 8). Asterisks depict significance (*, P < 0.05, **, P < 0.01). Error bars denote the SEM.

### Thalamic-cortical interactions are modulated by task context

We have shown above that for most recorded VPL-S1 pairs there were more feedforward interactions than feedback interactions during the first half (250 ms) of the stimulus-present. However, whereas these interactions were related exclusively to sensory information processing, but whether they were also influenced by the task’s context, remain unknown. To investigate this question, we applied our directionality analysis to a control task in which the monkey was passively stimulated by the same set of tactile stimuli but no perceptual report was required. Using task-balanced data sets (STAR Methods), we repeated some of the previous analysis for stimulus-present trials and stimulus-absent trials in both tasks, exploring potential differences in the percentage of directional interactions when the monkey was passively stimulated (Fig. 7). Based on the previous analysis of amplitude modulation (Fig. 6), we restricted our analysis during stimulus-present trials to near-threshold and supra-threshold amplitudes for feedforward interactions and supra-threshold interactions for feedback and bidirectional interactions.

**Figure 7.**
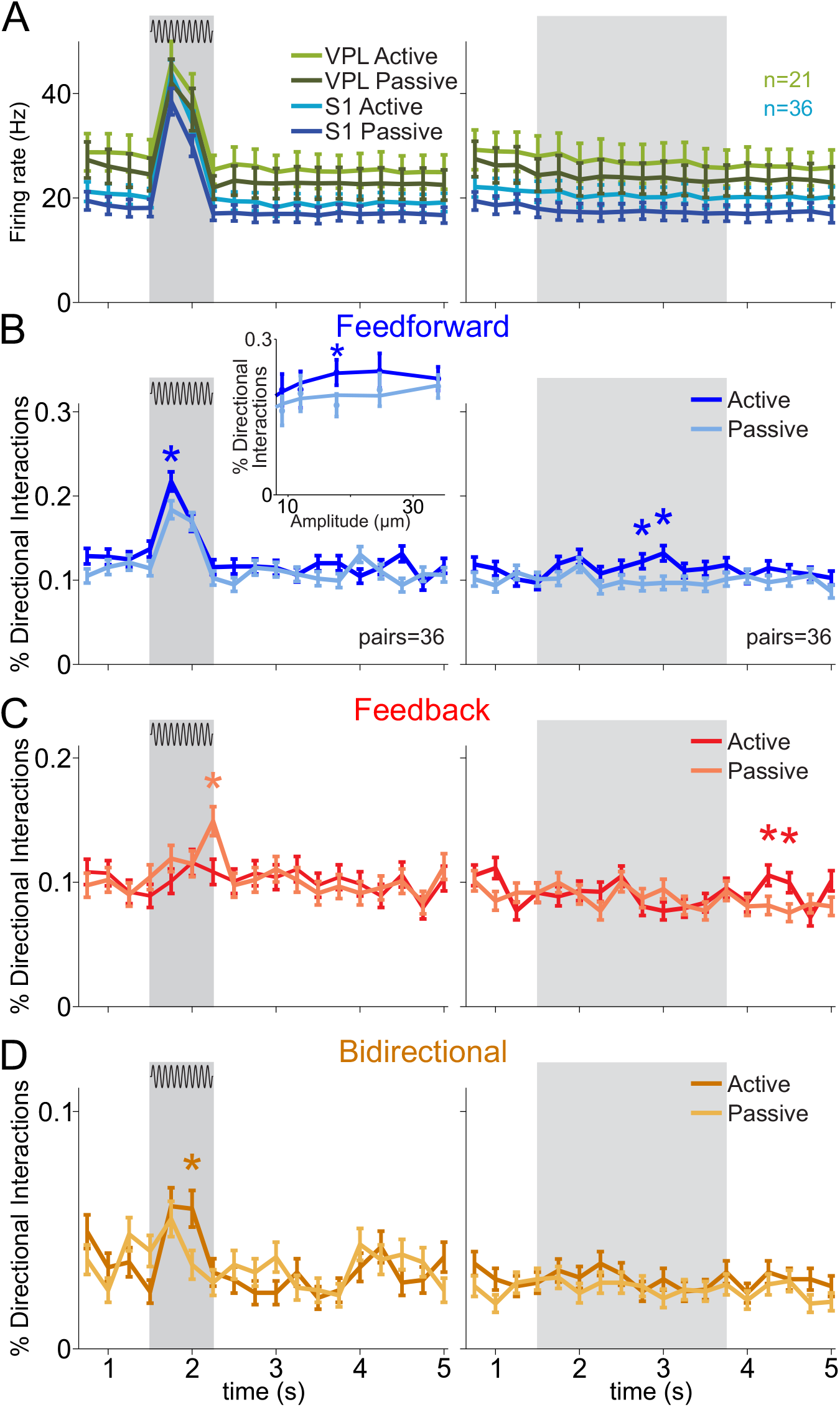
VPL-S1 interactions during passive stimulation. (A) Time course of the mean firing rates for VPL (n = 21) and S1 (n = 36) neurons. Neuron pairs (neuron pairs = 36) were simultaneously recorded during the detection task (active) and passive stimulation. For all panels, the left side depicts stimulus-present trials, whereas the right side depicts stimulus-absent trials. Dark and light colors depict data belonging to the active and passive conditions, respectively. (B) Time course of the percentage of feedforward interactions (neuron pairs = 36; trials = 1105 hits; trials = 1364 correct rejections). (C) Time course of the percentage of feedback interactions for the active and passive conditions (neuron pairs = 36; trials = 932 hits; trials = 1364 correct rejections). (D) Time course of the percentage of bidirectional interactions (neuron pairs = 36; trials = 932 hits; trials = 1364 correct rejections). Inset depicts the percentage of feedforward interactions as a function of the stimulus amplitude. Asterisks denote significance levels (*, P<0.05) associated with effect sizes (Cohen’s H) larger than 0.3.

We first revisited the firing rate of single neurons in VPL and S1 (Vázquez et al., 2012 and 2013) and found that the average (over neurons) firing rate of each area was not substantially altered during the timeline of the passive stimulation task (Fig. 7A). Consistent with this quantification, we thereafter analyzed the percentage of directional interactions during the passive condition over all neuron pairs that had been recorded in both tasks. In general, our analysis revealed task-specific variations at the level of directional interactions, which occurred around the stimulation window for stimulus-present trials (left panels, Figs. 7B, 7C and 7D). Besides that, significant feedforward differences were found during the PSW for stimulus-absent trials (right panel, Fig. 7B). As compared to the vibrotactile task, the arrival of the (supra-threshold) stimulus in the passive condition produced a lesser increase of bidirectional interactions (neuron pairs = 36; Fig. 7D left) during the second half of the stimulus window and a specific increase of feedback interactions during the first 250 ms of the post-stimulation window (neuron pairs=36; Fig. 7C left). Notably, feedforward interactions were significantly higher in the active than the passive condition, both during the stimulation period (left panel) and PSW (right panel). Thus, our findings show that the VPL-S1 interactions were sensitive to the task context. In particular, passive stimulation mitigated the number of bidirectional interactions during stimulus delivery and enhanced feedback interactions when the stimulus was no longer present.

### Feedforward interactions correlate with the animal’s task performance

An important question if whether the VPL-S1 interactions are modulated by the animal’s task performance. To further examine this question, we first analyzed the differences between hit and miss trials during stimulus-present condition (Fig. 8 left) and between correct rejections and false alarm trials during stimulus-absent condition (Fig. 8, right) across single neuron firing rates (Fig. 8A) and interactions types (Fig. 8B-D). In stimulus-present trials, we controlled for the effect of stimulus amplitudes by analyzing the difference between hits and misses at the near-threshold amplitude value (9 μm), where the number of samples was the most balanced between hit (trials = 443 hits in neuron pairs = 79) and miss responses (trials = 389 misses in neuron pairs = 79). In both experimental conditions, we controlled for a possible sample bias and experimental sessions effect, by using group-based permutation tests at the level of neuronal pairs (STAR Methods). We then outlined task intervals where the difference was significant and the effect size was larger than 0.2. Our analysis primarily revealed that the number of feedforward interactions during the first half of the stimulus period in stimulus-present trials was significantly larger (P < 0.05, H > 0.2) in hit trials (Fig. 8A, left). In contrast, our dataset did not unravel strong significant differences between correct rejections and false alarms during stimulus-absent trials (trials = 4188 correct rejections and false alarm trials = 933 in neuron pairs = 82). Overall, our results indicate that the amount of feedforward interactions during the first half of the stimulus window was correlated with the monkey’s behavior and could potentially predict the monkey’s performance in stimulus-present trials.

**Figure 8.**
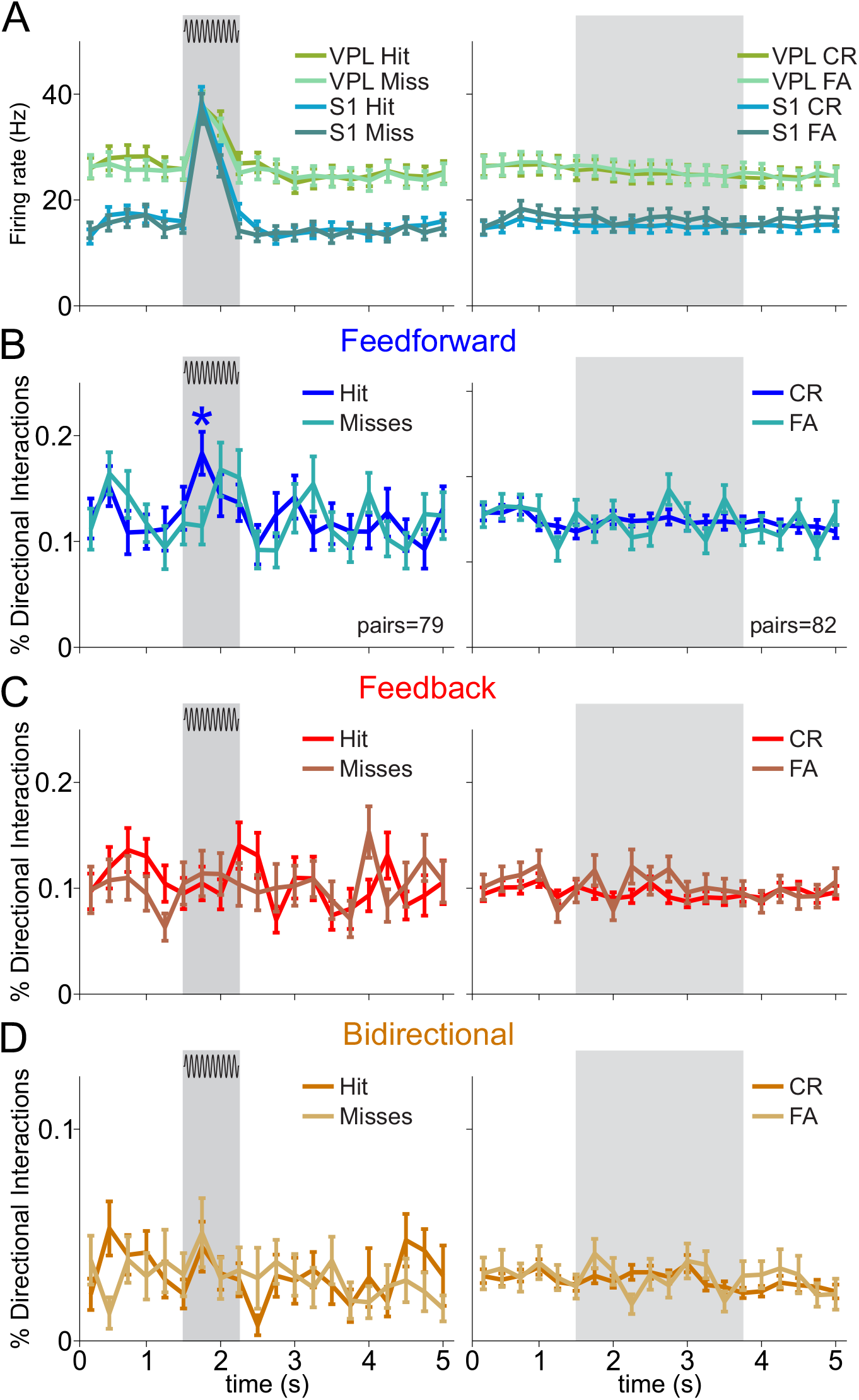
VPL-S1 interactions during correct and error trials. (A) Time course of the mean firing rate VPL (n = 48) and S1 (n = 70) neurons during stimulus present trials and for VPL (n=51) and S1 (n=73) neurons during stimulus-absent trials. Neuronal responses were separated according to the monkey’s behavioral output. Left panel represents hits and misses for the stimulus-present trials, whereas the right panel represents correct rejections and false alarms for stimulus-absent trials. Neurons and pairs were selected to have more than two trials at near-threshold amplitude for each condition. (B) Time course for the percentage of feedforward interactions. (C) Time course for the percentage of feedback interactions. (D) Time course for the percentage of bidirectional interactions. Left panels in B-D depict hits (dark colors) and misses (light colors) for the stimulus-present trials (neuron pairs = 79; trials = 443 hits; trials = 389 misses). Right panels in B-D depict correct rejections (dark colors) and false alarms (light colors) for the stimulus-absent trials (neuron pairs = 82; trials = 4188 correct rejections; trials = 933 false alarms). Asterisks denote significance levels (*, P<0.05) associated with effect sizes (Cohen’s H) larger than 0.3.

## DISCUSSION

Here, we sought to determine the functional roles of thalamo-cortical interactions in perception. This was done by measuring the directed information flow between simultaneously recorded VPL-S1 neuron pairs sharing the same cutaneous receptive field in monkeys performing a vibrotactile detection task. Notably, these interactions could not be merely explained by increases in the firing rates of neurons within VPL or S1 during the stimulus period. We found that the stimulus-presence elicited a significant increment on the number of feedforward (VPL→S1) and bidirectional (S1↔VPL) interactions. In contrast, pure feedback (S1→VPL) interactions remained practically unaltered during the stimulus-presence. Remarkably, increments in feedforward and bidirectional interactions were differently modulated as a function of the stimulus amplitude. Specifically, we found a linear relationship between feedforward interactions and stimulus amplitude, whereas bidirectional interactions emerged only with supra-threshold sensory inputs during task performance. Additionally, we identified that thalamic and cortical neurons involved in bidirectional interactions exhibited higher firing rates during stimulus-presence. Interestingly, the amount of feedforward interactions was correlated with the monkeys’ performance when they judged the presence of near-threshold stimuli. Also, during a passive stimulation task, when the monkeys were not required to report their percepts, feedforward interactions during the stimulus-presence were reduced. This suggests that feedforward interactions are modulated by the task-context. Finally, we compared thalamo-cortical interactions with those neurons that were recorded in the same area (S1-S1 and VPL-VPL) during the detection task. We found that unidirectional and bidirectional S1-S1 interactions were modulated during stimulus presence, but not in the VPL-VPL interactions. We discuss these findings below.

Previous studies aimed to assess the directionality of the thalamo-cortical interactions. Most of them have inferred directionality on the frequency domain (Bastos, 2014; FitzGerald et al., 2013; Haegens et al., 2014; Wang et al., 2007). Examples of such metrics are partial directed coherence between accumulated single-unit activity in the ventral posterior medial (VPM) thalamic nucleus and S1 during a nociceptive experiment (Wang et al., 2007), and the phase slope index between the local field potentials (LFP) of the lateral geniculate nucleus (LGN) and visual cortex (V1) during a visual task (Bastos, 2014). Also, in a previous study with the same detection task used here, Haegens and collaborators (2014) used spike-field coherence (SFC) to identify oscillatory synchronization between VPL and S1. The authors found that the magnitude of SFC was increased in the beta band during the stimulus period and was higher in the VPL→S1 direction than in the S1→VPL direction.

In the current study, we extended these observations to the temporal domain, at the level of single-neuron interactions. To achieve this, we analyzed directed information flow between VPL and S1 neuronal spike trains with a nonlinear measure (directed information; Massey, 1990; Jiao et al., 2013; Tauste Campo et. al, 2015). Indeed, we decomposed single-neuron interactions into three types: feedforward, feedback and bidirectional, and explored their temporal evolution from the pre-stimulus, stimulus up to the post-stimulus intervals of the detection task. In particular, the lack of feedback modulation during stimulation period, suggests that SFC increment in the S1→VPL direction (Haegens et al., 2014) could be associated with increments in bidirectional interactions. Our analytical approach involved the use of nonparametric significance test (Gilson et al., 2017) and the choice of the percentage of significant interactions as a relevant connectivity metric (Tauste Campo et al, 2015). Both choices were critical to detect nonlinear interactions in a way that was only weakly dependent on the firing rate. As a result, we were able to analyze neuronal data in a dimension that was quasi-orthogonal to the firing rates (Vázquez et al., 2012 and 2013), while still showing rich stimulus modulations. Besides, it is important to mention the differences between the directed information used here with the widespread measure of noise correlation (Ecker et al., 2014; Kohn et al., 2016). Noise correlation averages across trials under the same stimulus condition, correlating fluctuations in firing rate of two neurons. In contrast, directed information quantifies, in a single trial and for any given time, the information that the recent past and present spike train of a given neuron has about the present spike train of the other, simultaneously recorded, neuron.

Notably, feedforward VPL-S1 interactions were modulated during the stimulus period. Further, the modulation was stronger 250 ms immediately after the stimulus onset. These findings are congruent with the adaptation of feedforward interactions over the stimulus period. Indeed, previous studies have shown that thalamic and cortical neurons become adapted to tactile stimuli after several pulses (Khatri et al., 2004, Vázquez et al., 2012 and 2013). Moreover, it has been reported that this adaptation changes the neural code of cortical neurons and thalamic synchrony from a detection to a discrimination modality (Wang et al., 2010). As stated above, our directionality measure is poorly correlated with differences in firing rate of VPL and S1 neurons. Therefore, our results are congruent with the adaptive coding paradigm (Wang et al., 2010), by showing that feedforward interactions exhibit the strongest modulation during the first part of the stimulus-presence. In sum, we hypothesize that feedforward adaptation reflects an internal mechanism that prioritizes the information contained within the first pulses of the sinusoidal stimulus to further transmit this information to the cortex and therefore, for stimulus detection during this task.

According to the standard classification of thalamic sensory nuclei (Sherman, 2016), VPL is considered a first-order relay nucleus that mainly transmits afferent information to S1 in the presence of cortical feedback (Ahissar and Oram, 2015; Blomquist et al., 2009; Crandall et al., 2015). Our results support this view by showing that feedforward interactions alone were primarily boosted during the first 250 ms immediately after the stimulus onset. The increment in feedforward interactions was concentrated at an interaction delay of 8 ms, which is consistent with previous literature (Briggs and Usrey, 2008). Moreover, these increments were reduced during incorrect trials and, to a lesser extent, during the passive control condition. In particular, the relationship between feedforward interactions and the monkey’s performance during stimulus-present trials is of special novelty. In previous related studies, this correlation did not arise neither in spiking activity (Vázquez et al., 2013) nor in the oscillatory activity (Haegens et al., 2014). This result suggests that perceptual detection might be related to the amount of stimulus information flowing from VPL to S1. In contrast, pure feedback interactions did not display substantial task modulations. We hypothesize that feedback interactions might be relevant for perceptual detection, but are generally less frequent than feedforward interactions. Hence, larger samples of simultaneous VPL-S1 neuron pairs might be needed to explore their real functional relevance in the detection task. On the other hand, different cortical areas might show more prominent feedback cortico-thalamic interactions, perhaps related to not only facilitating the gating of the stimulus, but also some other cognitive aspects not detected in this task. But, see below.

Our study revealed that feedback interactions could take place almost simultaneously with the feedforward interactions, suggesting the emergence of a thalamo-cortical loop, denominated here as bidirectional interactions. Notably, our analysis identified three relevant singularities of these bidirectional interactions. First, they involved larger directed information values that occurred mostly at 0 ms delay. Second, the amount of bidirectional interactions was (weakly) modulated by the stimulus amplitude: they appeared only with supra-threshold stimulus amplitudes. Third, thalamo-cortical neurons involved in bidirectional loops, exhibited higher firing rates during stimulus period. Remarkably, bidirectional interactions mostly increased 250 ms immediately after the stimulus onset. We speculate that this increase might be explained by three possible causes. First, it might be due to an increment of inputs from indirect pathways through cortical neurons from layers 4 and 6 of S1, very likely via the reticular nucleus (Alitto and Usrey, 2003). Second, it could reflect the correlated response of a tactile population code across VPL and S1 neurons (Kohn et al., 2016). Third, the highly informative 0 ms delay bidirectional interaction could be related to a coordinate mechanism for phase locking the neuronal activity between VPL and S1 (binding in synchrony, Singer, 1999). Intriguingly, the second highest increase in bidirectional interactions occurred again at 8 ms immediately after the stimulus onset. This interaction delay might be associated with reciprocal connections, which have been described as a fast and reliable way to process stimulus information, as shown between the lateral geniculate nucleus and V1 (Briggs and Usrey, 2008; Diao et al., 2017).

In several sessions we could simultaneously record pairs of neurons in the VPL or in the S1, while the monkeys performed the task. This allowed the analysis of the intra-thalamus (VPL-VPL) and intra-cortical (S1-S1) interactions. Notice that the neurons that were recorded in each area shared the same cutaneous receptive field. Importantly, we did not identify significant modulation on intra-thalamus interactions. Even if the number of VPL-VPL pairs is small (it is challenging to simultaneously record VPL neurons sharing their small receptive fields), our results support the idea that VPL neurons are more involved in relaying sensory information to S1, with few interactions between them (Ahissar and Oram, 2015; Sherman and Guillery, 2006). However, to further test the role of the VPL-VPL interactions, future experiments must increase the number of simultaneously recorded thalamic neuron pairs. In contrast, unidirectional and bidirectional intra-cortical (S1-S1) interactions exhibited modulation during the stimulus-presence, which was more sustained over the entire stimulus period than in VPL-S1 pairs. This agrees with previous studies suggesting that the neural interaction in S1 could contribute to the stimulus processing (Reed et al., 2008).

In brief, the present study employed a refined methodology to support that the increment of VPL→S1 neuronal interactions (as opposed to S1→VPL) during the stimulus period, reflects the effective transmission of tactile information from thalamus to cortex. Moreover, it suggests that the amount of feedforward interactions occurring during the first 250 ms of stimulation correlate with the monkey’s stimulus perception and with the behavioral task performance. Notoriously, the bidirectional (VPL↔S1) interactions constitute a novel and coordinated mechanism to transmit information between these two areas. Our results, therefore, contribute to understanding the directed information flow between the VPL and S1 during the detection of a tactile stimulus. Finally, the minimalistic approach used here; that is, the simultaneous recording of VPL-S1 neuron pairs sharing the same receptive field, together with the estimation of the directed information flow, could be used to not only investigate the thalamo-cortical interactions during perception, but also across other brain areas in this and in other behavioral tasks.

## ACKNOWLEDGMENTS

This work was supported in part by the Dirección General de Asuntos del Personal Académico de la Universidad Nacional Autónoma de México (PAPIIT-IN202716) and Consejo Nacional de Ciencia y Tecnología (CONACYT-240892) (R.R.). Support for this work was provided by European project FP7-ICT BrainScales (to G.D. and M.M.-G). In addition, G.D. was supported by the European Research Council Advanced Grant: DYSTRUCTURE (n. 295129), by the Spanish Research Project SAF2010-16085. YVZ was supported by a Pew Latin American Fellowship and a Charles H. Revson Biomedical Science Fellowship.

## SUPPLEMENTAL INFORMATION

Supplemental Information includes four figures and can be found with this article.

## AUTHOR CONTRIBUTIONS

A.T.C., Y.V., M.A., A.Z., R.R.-P., G.D., and R.R. contributed to different aspects of the study, including design and performance of research, data analysis, and writing. R.R. supervised all stages of the study.

## STAR METHODS

### METHOD DETAILS

#### Detection task

Stimuli were delivered to the skin of the distal segment of one digit of the restrained hand, via a computer-controlled stimulator (BME Systems, MD; 2-mm round tip). The initial probe indentation was 500 μm. Vibrotactile stimuli consisted of trains of 20 Hz mechanical sinusoids (20 ms duration each sinusoid), with amplitudes of 1-34 μm (Fig. 1A). These were interleaved with an equal number of trials where no mechanical vibrations were delivered to the skin (amplitude = 0). A trial began when the probe tip (PD) indented the skin of one fingertip of the restrained, right hand, upon which the monkey placed its free, left hand on an immovable key (KD). After a variable pre-stimulus period (1.5-3 s), a vibrotactile stimulus could be presented or not (0.5 s). After a fixed delay period (3 s), the stimulator probe was lifted off from the skin (PU), indicating to the monkey that it could initiate the response movement (KU) to one of two buttons (PB). The button pressed indicated whether or not the monkey felt the stimulus (henceforth referred as ‘yes’ and ‘no’ responses, respectively). They were rewarded with a drop of liquid for correct responses. Psychometric detection curves were obtained by plotting the proportion of yes responses as a function of the stimulus amplitude (left panel of Fig. 1B). Depending on whether the stimulus was present or absent and on the behavioral response, the trial outcome was classified as hit, miss, false alarm or correct rejection (right pane of Fig. IB). Monkeys were handled according to the institutional standards of the National Institutes of Health and Society for Neuroscience. All protocols were approved by the Institutional Animal Care and Use Committee of the Instituto de Fisiología Celular of the National Autonomous University of Mexico (UNAM).

In addition to the experimental condition described above, the animals also performed a passive control task (referred as passive condition) during which the stimulus was present or absent, but no response was required (Vázquez et al., 2012). Monkeys were rewarded randomly during the occurrence of the passive condition.

Under this situation, sensory information enters or not to the somatosensory system, but no decision and perceptual report is required to obtain a reward.

#### Recordings

Neuronal recordings were obtained by using two arrays, each with seven independent, movable microelectrodes (2 −3 MΩ; Hernández et al., 2008; Romo et al., 1999). One Array was inserted into S1 (cyan spot on the figurine of left panel of Fig. 1C), in the cutaneous representation of the fingers (areas 1 or 3b; middle panel of Fig. 1C). The other array was located lateral and posterior to the hand’s representation (green spot on the figurine of left panel of Fig. 1C) in a way that allowed us to lower the microelectrodes to the cutaneous representation of the fingers in the VPL of the somatosensory thalamus (right panel of Fig. 1C). Recordings were performed contralateral to the stimulated hand (right) and ipsilateral to the responding hand (right). Each recording began with a mapping session to find the cutaneous representation of the fingers in VPL. Subsequently, we mapped neurons in S1 sharing receptive fields with the neurons of VPL (Fig. 1D). All recorded neurons had small cutaneous receptive fields with quickly (QA, VPL n = 65, S1 n=71) or slowly adapting (SA, VPL n = 9, S1 n=4) properties. Locations of the electrode penetrations in VPL and S1 were confirmed with standard histological techniques. The neuronal signal of each microelectrode was sampled at 30 kHz and spikes were sorted online. A more extensive description of the task and recording procedure can be found in previous publications (Hernández et al., 2008; Vázquez et al., 2012).

Here, we report data from multiple recording sessions during which spikes were obtained. For the experimental condition, we recorded 47 sessions with 120−140 trials per session (53 neurons in VPL, 75 neurons in S1, 84 VPL-S1 pairs). For the passive control condition, we obtained 21 sessions with 70-140 trials (21 neurons in VPL, 36 neurons in S1). We performed a fully balanced comparative analysis between the original and control task recordings to controlled for statistic bias. To do so, we only considered VPL-S1 neuron pairs that were recorded in both experimental conditions. In addition, for each of these pairs, we performed correct trial subsampling to obtain the same type and number of amplitude classes recorded in the vibrotactile detection and the passive stimulation task. As a result, the pairing pre-processing yielded 36 VPL-S1 pairs that were recorded in both conditions: 1307 stimulus-present trials and 1364 stimulus-absent trials that could be used for unbiased statistical comparison. In the comparative analysis, we removed the first two intervals of both tasks due to the presence of signal artifacts in the passive condition. Hence, the analysis was restricted to the sub-period 0.5 − 5s in both tasks, which was in turn distributed over 18 non-overlapping task intervals (0.25s).

### QUANTIFICATION AND STATISTICAL ANALYSIS

#### Single-trial directionality analysis

We used custom-built MATLAB codes to analyze the data. The directionality analysis presented here is a refinement of our previous method to analyze spike-train directional correlations (Tauste Campo et al, 2015). We estimated neural directional correlations between every neuron pair within a population using a Bayesian estimator of the directed information (*DI*) (Massey, 1990) between a pair of discrete time series that were assumed to be generated according to a Markovian process. In more specific terms, for a pair time series (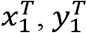) of length *T*, where 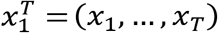 and 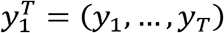, a time delay *δ*≥0, and Markovian orders equal to *D_1_* > 0 and *D*_2_ > 0, respectively, the *DI* between the stationary processes of 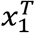 and 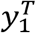, i.e., *X* and *Y*, is estimated through the formula:

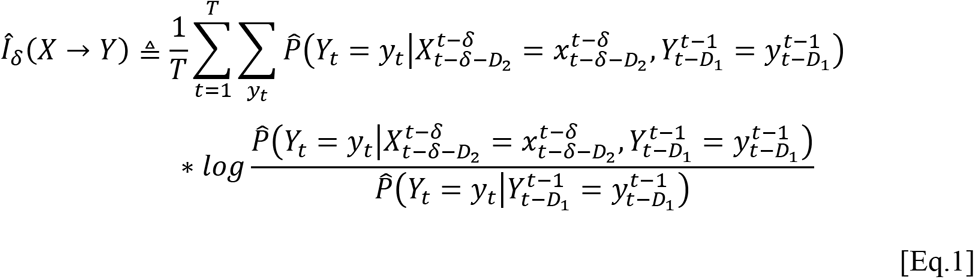

Where, the probability distribution of (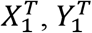) is estimated using the context-tree weighting algorithm (CTW, Jiao et al 2013). Equation 1 quantifies the information in that the past of 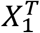 at delay *δ*, i.e., 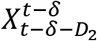, has about the present of 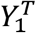, i.e., *Y_t_*, given the most recent part of 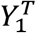, i.e., 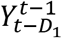. This estimator is consistent as long as (*X, Y*) the two neuronal time series, form a jointly stationary irreducible aperiodic finite-alphabet Markov process whose order does not exceed the prescribed maximum depth in the CTW algorithm (Jiao et al., 2013). Prior to estimating the *DI*, we preprocessed our data as follows. For a single trial, we first binarized spike-train trials using bins of 1ms (mapping 1 to each bin with at least one spike and 0, otherwise). Second, in stimulus-present trials, we removed the variable-time pre-stimulus period in every trial and aligned all trials to the stimulus onset time. In contrast, in stimulus-absent trials, we aligned the trials to the probe down event (PD). We then divided each trial time series into twenty non-overlapping task intervals of 0.25s (250 bins). For each neuron and segment, the spike train was assumed to be generated by a random process that satisfied the estimator requirements with a maximum memory of 2 ms (*D*_1_ = *D*_2_ = 2 bins) both for the joint and the marginal spike-train processes. Under this assumption, it can be easily checked that the *DI* is asymptotically equivalent to the transfer entropy (Schreiber, 2000) in the limit of the time-series length. To assess that neurons were able to express minimal information through their spike-train responses, we ran the estimator of the entropy (a particular case of the *DI* estimator) for each neuron and task interval time series with maximum memory, *D* = 2. This step removed spike trains with zero or small number of spikes. Finally, among those neurons that had a significant entropy value, we ran the *DI* estimator (Tauste Campo et. al, 2015) over all possible pairs and time delays 0, 2, 4, 6, 8, 10, 12, 14, 16, 18, 20 ms.

To assess statistical significance for both entropy and *DI* estimations, we used a Monte-Carlo permutation test (Ernst, 2004). In this test, the original (i.e., non-permuted) estimation was compared with the tail of a distribution obtained by performing 20 equally spaced circular shifts of the target spike train *Y^T^* (*α*=5%) and computed the corresponding P-value. We dealt with the multiple test problem over delays in the *DI* estimation by using the maximum *DI* over all preselected delays as a test statistic. As a result, the test provides three outputs: the interaction significance (0/1), the interaction strength (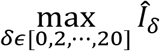), and the maximizing delay *δ̂*. Then, every time-vaying percentage of directional interactions (feedforward, feedback and bidirectional) plotted in Figs. 3-8 was obtained by dividing the number of significant interactions of the corresponding type at a given task interval over all neuron pairs and trials (Figs. 3-7), and over all trials per neuron pair and then averaging the percentages over all neuron pairs (Fig. 8).

#### Statistical analysis

Main results illustrated in Figs. 3-8 were obtained by comparing time-varying percentages of significant interactions at each task interval. Comparisons were of two types: paired and unpaired. Paired comparisons appeared in the comparison between the VPL→S1 and VPL→S1 percentages (Fig. S1), the percentages over neuron pairs in different conditions (Fig. S2 and S3), the average firing rate over neurons holding directional interactions (Fig 4), the percentages between the original and control task (Fig. 7) and the percentages over neuron pairs for correct and error trials (Fig. 8). Unpaired comparisons appeared when assessing the stimulus-driven change in the percentage of directional interactions (Fig. 3 and 6). In both comparisons, we used non-parametric tests for correlated samples (Winkler et al., 2014) using statistics based on Cohen’s effect size (Cohen’s h; Cohen, 1988) that measured the distance between proportions. The use of this statistic allows to straightforwardly quantify the size of any significant effect by comparing its value with standarized thresholds (*H* = 0.2, small effect size; *H* = 0.5, medium effect size; *H* = 0.8 large effect size), thus avoiding sample size biases. For any unpaired comparison between proportion *p*_1_ and *p*_2_, we used the original Cohen’s h measure:

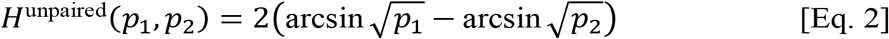

For paired comparison, we proposed the following paired version of Cohen’ h:

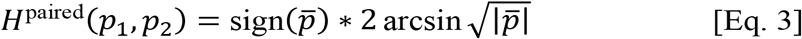

where 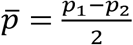.

Non-parametric tests for correlated samples were performed through 1000 group-based permutations (Winkler et al., 2014) where groups were defined to be either single trials (Figs. 3-7) or neuronal pairs (Fig. 8) and group sample sizes were maintained in each permutation. Thus, our analysis avoided introducing any statistical bias to the sampled reference distribution. In the main study, comparisons were sequentially performed over all task intervals (*N* = 20). To correct for test multiplicity, we applied the Holm-Bonferroni procedure (Holm, 1979), which provided a significance threshold that controlled the Familywise Error Rate (FWER) at a significance level (*α* = 0.05). In the remaining tests performed at different amplitude values or neuron pairs, multiplicity was not corrected for lack of sufficiently large sample sizes.

In Fig. 6B we applied single-trial and average-trial correlation measures to quantify the non-parametric correlation (Spearman’s rho) between percentages of interactions and stimulus amplitudes. Single-trial correlation values were obtained by correlating the trial-based binary vector associated to each interaction type against their corresponding amplitudes. Average-trial correlations were obtained by first computing the overall percentage of interactions at each amplitude and then by correlating these percentages against all amplitude values.

---

**Figure S1 related to Figure 3.**
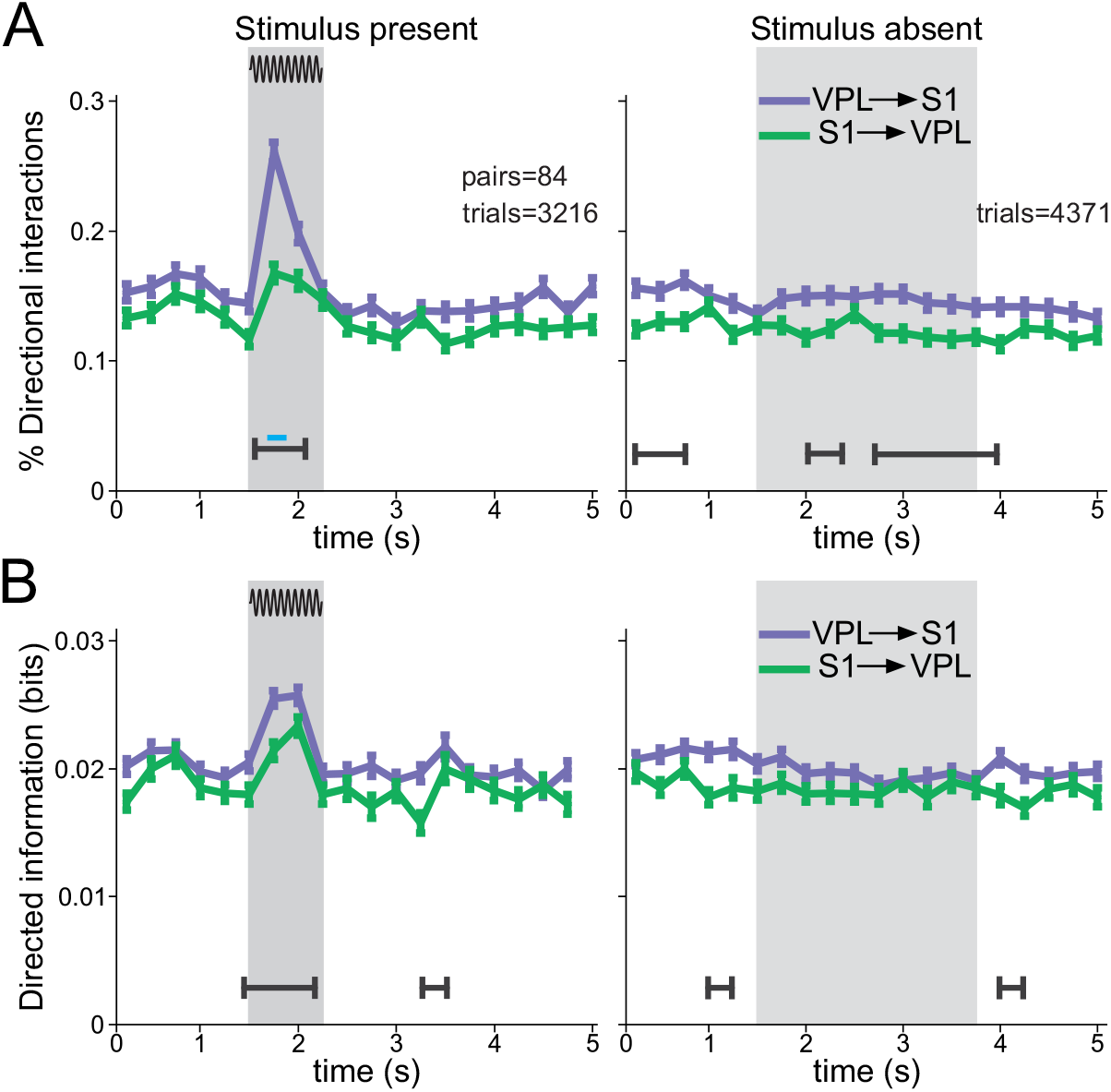
Time course of the task during directional interactions and directed information between VPL→S1 (purple) and S1→VPL (green) in stimulus-present (left panel, trials = 3216 hit; neuron pairs = 84) and stimulus-absent trials (right panel, trials = 4371 correct rejections; neuron pairs = 84). In all figures, grey boxes depict the stimulation period for the stimulus-present trials, and the possible window of stimulation (PWS) for the stimulus-absent trials). Error bars denote the SEM (standard error of the mean). (A) Percentage of directional interactions. Black lines depict intervals for which the difference between directional interactions was significantly different (non-parametric unpaired test P < 0.05, multiple-test corrected; effect size > 0.3). Blue lines depict significant intervals for which the effect size was larger than 0.5. (B) Time course of directed information for VPL→ S1 (purple) and S1→ VPL (green) interactions. Black lines depict intervals for which the difference between directions was significantly different (non-parametric unpaired test P < 0.05, multiple-test corrected; effect size > 0.15).

**Figure S2 related to Figure 3.**
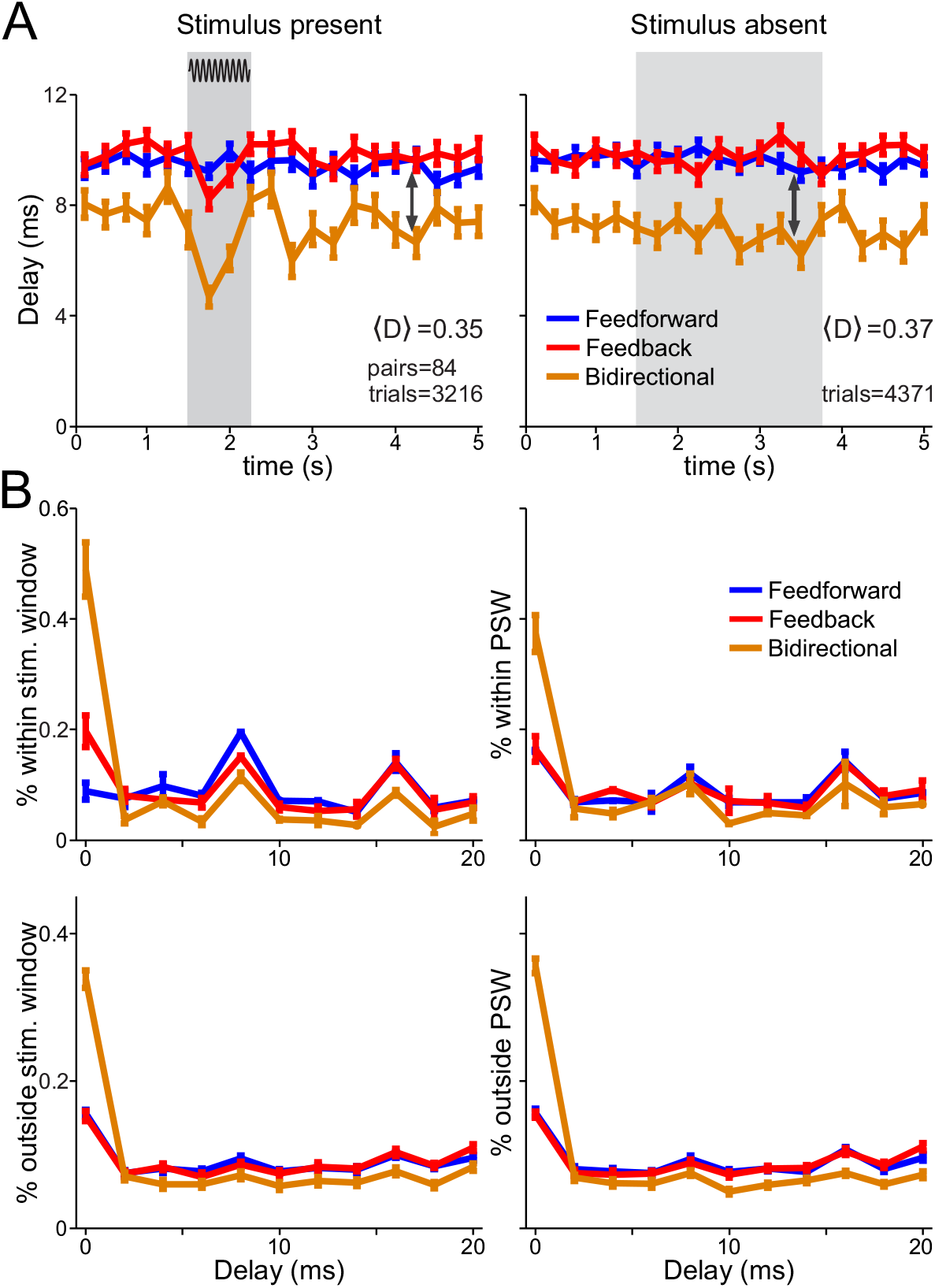
Mean delay for feedforward, feedback and bidirectional interactions. (A) Time course of the task for the mean delays occurring during feedforward, feedback and bidirectional interactions for stimulus present (left, trials=3216 hits; neuron pairs = 84) and stimulus absent (right, trials=4371 correct rejections; neuron pairs = 84) trials. 〈D〉 Indicates the average value of the effect size (over task intervals and interaction types) between feedforward, feedback, and bidirectional interactions. (B) Percentage of delays within (upper panel) or outside (lower panel) the stimulation or PSW window for stimulus present (left) and stimulus absent (right) trials.

**Figure S3 related to Figure 3.**
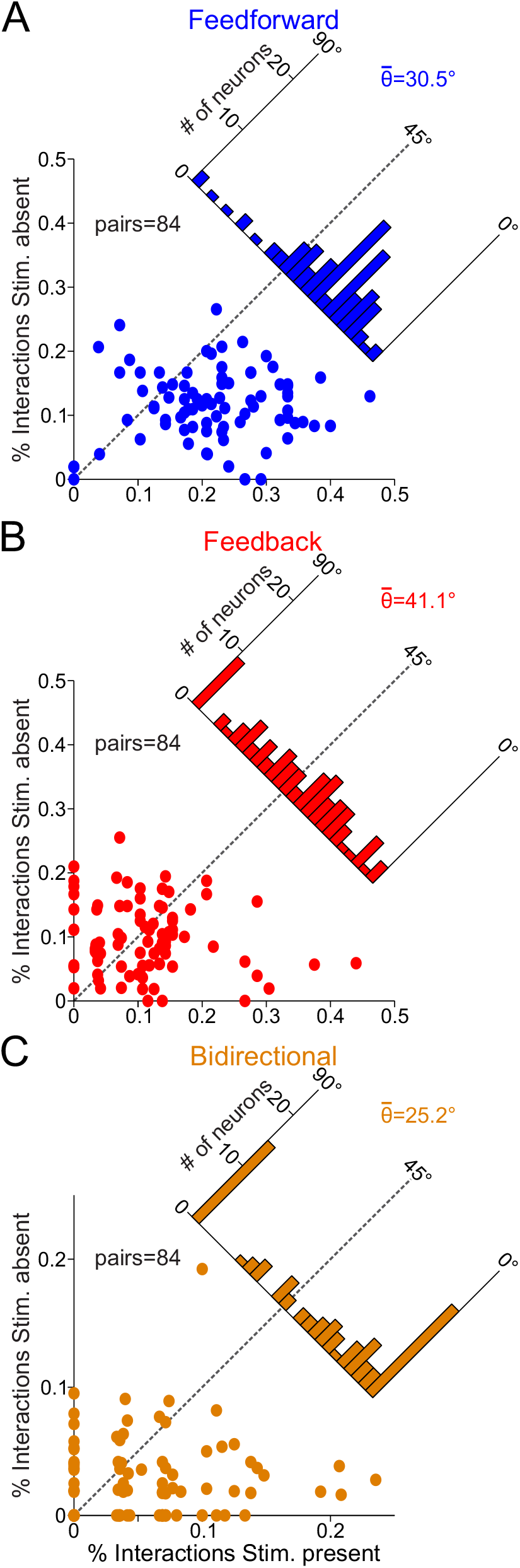
Percentage of (A) feedforward, (B) feedback, and (C) bidirectional interactions per neuron pair (trials = 3216 hits; trials = 4371 correct rejections; neuron pairs = 84) during the first interval (250 ms) of the stimulus period in stimulus-present trials vs. the first interval (250 ms) of the PSW in stimulus-absent trials. In each panel the insets depict the histograms of the angular deviation between stimulus-absent and stimulus-present trials over all neuronal pairs and indicates its median. The percentage for all three types of interactions was higher during the stimulus-present trials (θ < 45°) with feedforward (θ = 30.5°) and bidirectional (θ = 25.2°) interactions exhibiting larger differences across conditions than feedback interactions (θ = 41.1°).

**Figure S4 related to Figure 4.**
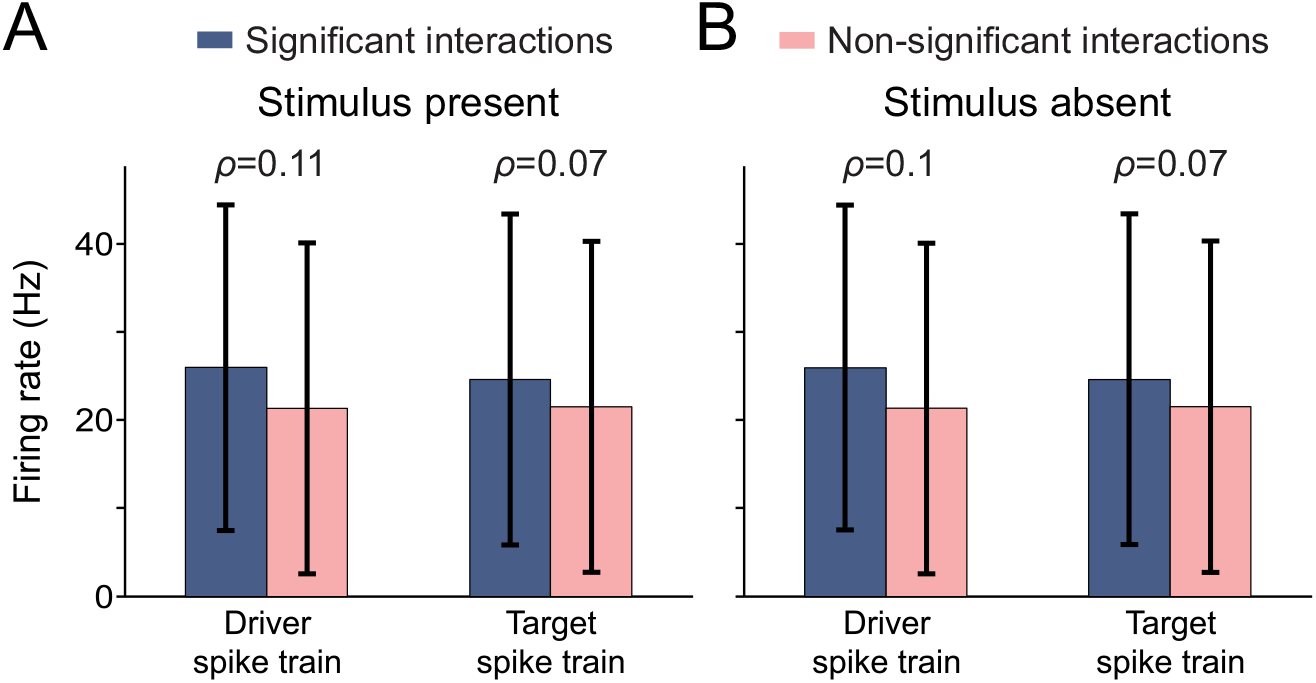
Mean firing rate for driver and target neurons having significant and non-significant directional interactions across the VPL→S1 and S1→VPL neuron pairs (84). (A) Mean firing rate and standard deviation for driver and target neurons for the stimulus-present trials. Blue depicts firing rates related to significant interactions (significant intervals = 29074; trials = 3216 hits). Pink depicts firing rates related to non-significant interactions (non-significant intervals = 177286). The value of ρ indicates the Spearman correlation value obtained from correlating the firing rate of the driver/target with the existence of incoming/outgoing directional interactions. (B) Same as in (A) but for the stimulus-absent trials (significant intervals = 28253; non-significant intervals = 178027; trials = 4371 correct rejections).

